# Regulatory plasticity within a complex cytokine-sensing mammary enhancer during lactation

**DOI:** 10.1101/2020.06.04.134429

**Authors:** Hye Kyung Lee, Chengyu Liu, Lothar Hennighausen

**Affiliations:** Laboratory of Genetics and Physiology, National Institute of Diabetes and Digestive and Kidney Diseases, US National Institutes of Health, Bethesda, Maryland 20892, USA; Transgenic Core, National Heart, Lung, and Blood Institute, US National Institutes of Health, Bethesda, Maryland 20892, USA

## Abstract

Enhancers are transcription factor platforms that synergize with promoters to activate gene expression up to several-thousand-fold. While genome-wide structural studies are used to predict enhancers, the *in vivo* significance is less clear. Specifically, the biological importance of individual transcription factors within enhancer complexes remains to be understood. Here we investigate the structural and biological importance of individual transcription factor binding sites and redundancy among transcription components within a complex enhancer *in vivo*. The *Csn1s2b* gene is expressed exclusively in mammary tissue and activated several thousand-fold during pregnancy and lactation. Using ChIP-seq we identified a complex lactation-specific candidate enhancer that binds multiple transcription factors and coincides with activating histone marks. Using experimental mouse genetics, we determined that deletion of canonical binding motifs for the transcription factors NFIB and STAT5, individually and combined, had a limited biological impact. Loss of these sites led to a shift of transcription factor binding to juxtaposed sites, suggesting exceptional plasticity that does not require direct protein-DNA interactions. Additional deletions revealed the critical importance of a non-canonical STAT5 binding site for enhancer activity. Our data also suggest that enhancer RNAs are not required for the activity of this specific enhancer. While ChIP-seq experiments predicted an additional candidate intronic enhancer, its deletion did not adversely affect gene expression, emphasizing the limited biological information provided by structural data. Our study provides comprehensive insight into the anatomy and biology of a composite mammary enhancer that activates its target gene several hundred-fold during lactation.

## Introduction

Enhancers are transcription component platforms that control the location, timing and intensity of gene expression^1,2^. While current approaches, such as the ChIP-seq and physical contact studies, are useful in identifying candidate enhancers, their biological predictions are limited and validation through genetic experiments is needed. Candidate enhancers are commonly occupied by multiple transcription factors (TFs) that might bind through their respective DNA recognition motifs or indirectly through other proteins. Since experimental genetic studies generally ablate the entire enhancer, the structural and functional contribution of individual TFs is poorly understood. Specifically, the functional plasticity and possible compensation of individual enhancer components is not understood.

Several hundred genes are uniquely expressed in mammary tissue and activated by pregnancy and lactation hormones through the tyrosine kinase JAK2 and the transcription factor Signal Transducer and Activator of Transcription (STAT) 5^3–5^. While most STAT5 target genes are highly induced during pregnancy and to a lesser extent during lactation^6^, the activation of the *Csn1s2b* gene^7^ occurs preferentially during lactation^8^. Previously, we identified enhancer structures that were preferentially established during lactating^8^. ChIP-seq profiles for STAT5 and H3K27ac and other mammary-enriched TFs suggested the presence of highly complex mammary enhancers^8^. Although most of these enhancers appear to depend on STAT5 as the anchor for the establishment of larger protein complexes, the stage-specific establishment of enhancers remains to be understood. It is not known why seemingly structurally identical enhancer can be activated by pregnancy hormones either during pregnancy or lactation.

Here, we investigated a potential synergy between the prolactin-induced TF STAT5 and the mammary-enriched NFIB in the establishment of a lactation-specific enhancer that activates gene expression in mammary tissue several hundred-fold. For this we investigated the enhancer landscape during pregnancy and lactation and introduced specific mutations in the mouse genome. This permitted us to explore the contributions and significance of individual TFs in the establishment of a functional enhancer. Specifically, we focus on the *Csn1s2b* enhancer, which contains several distinct regulatory elements bound by the TFs STAT5A, NFIB and MED1 that control gene expression during lactation^8^.

## Results

### The *Csn1s2b* enhancer is activated in mammary tissue during lactation

*Csn1s2b* is a member of the casein (*Csn)* family that is expressed exclusively in mammary tissue under the control of pregnancy and lactation hormones (Supplementary Table 1). While four out of the five casein genes are highly induced during pregnancy, *Csn1s2b* is activated preferentially and up to several-hundred-fold during lactation, suggesting the presence of distinct regulatory elements. The juxtaposed *Csn1s2a* and *Csn1s2b* genes, which arose by gene-duplication prior to the split of eutherian mammals^9,10^, are subject to different regulation. While *Csn1s2b* expression increase more than 250-fold between day 1 of lactation (L1) and day 10 (L10), *Csn1s2a* expression increased approximately 6-fold (Fig. 1a) suggesting the presence of a unique enhancer that differentially respond to lactation hormones, with prolactin the most prominent one. Digging deeper, we used ChIP-seq profiling for transcription factors and histone marks to identify candidate enhancers (Fig. 1b-d and Supplementary Fig. 1a). Binding of the prolactin-activated STAT5 was detected at three sites upstream of the *Csn1s2a* gene and two sites at the *Csn1s2b* gene (Fig. 1b). The most proximal STAT5 binding at the two genes was in close proximity of the TSS, suggesting that they could be part of a combined promoter-enhancer structure. While maximum STAT5 binding at the *Csn1s2a* sites is observed at day 18 of pregnancy (p18) and remains high throughout lactation, it is preferentially established at *Csn1s2b* gene between L1 and L10 (Fig. 1b and Supplementary Fig. 1a-b). Thus, seemingly identical TF binding sites respond differently during pregnancy and lactation. Pol II loading and H3K4me3 coverage at the two loci also reflects the differential expression of the two genes (Fig. 1b-d).

**Fig. 1.**
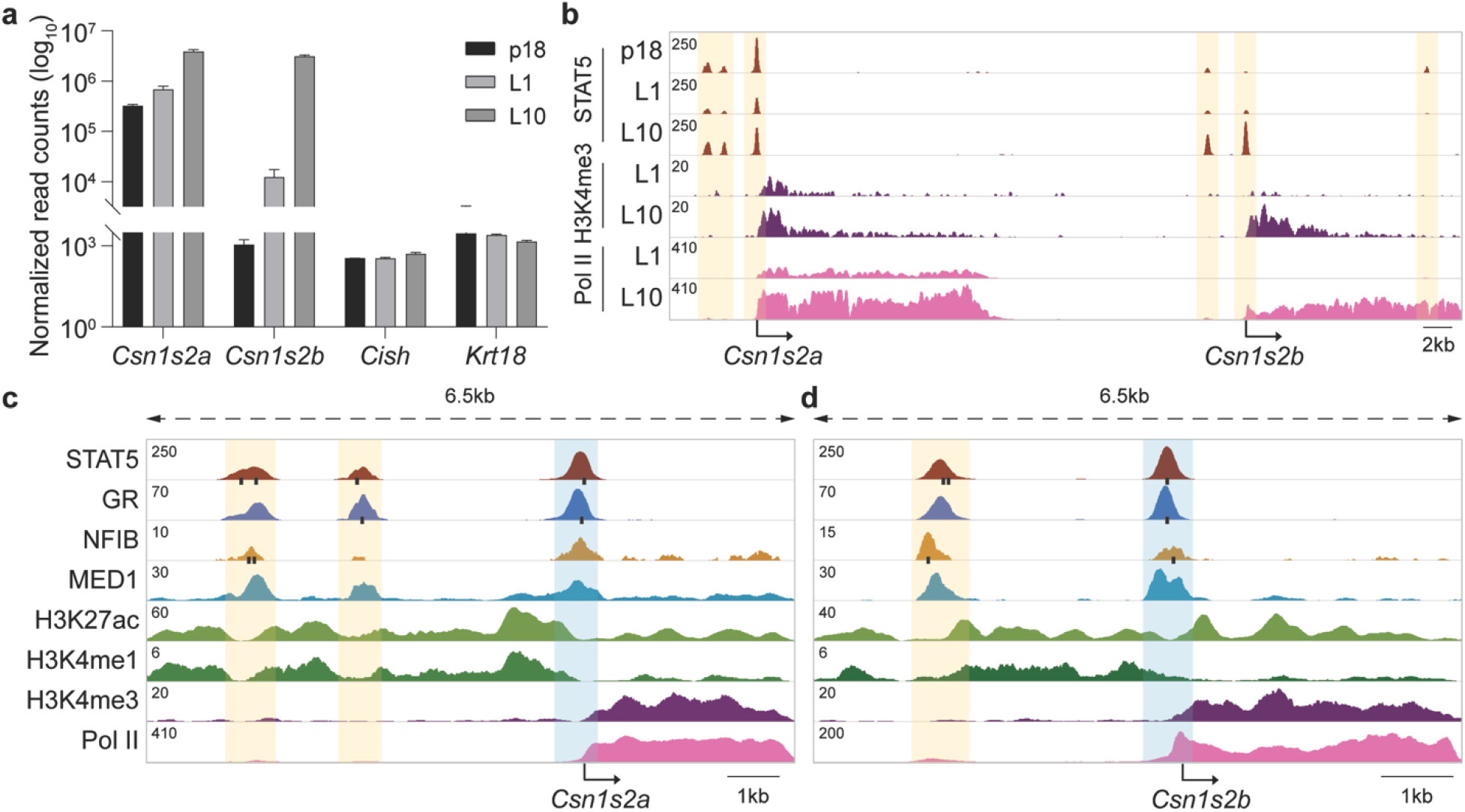
Structure and function of the *Csn1s2a/b* locus. **a** mRNA levels of both genes were measured by RNA-seq at day 18 of pregnancy (p18), day 1 of lactation (L1) and L10 (*n* = 3 or 4). The *Cish* and *Krt18* genes served as controls. **b** ChIP-seq data for transcription factors (TFs) and histone markers provided structural information of the *Csn1s2a*-*Csn1s2b* locus during pregnancy and lactation. The orange shades indicate putative regulatory elements. The *Cish* locus served as control (Supplementary Fig. 1). DE, distal enhancer; IE, intronic enhancer. **c-d** TF binding and activating histone marks at the *Csn1s2a* (c) and *Csn1s2b* (d) locus at L10. The black bars indicate the binding motifs of STAT5, GR and NFIB. The orange and blue shades indicate putative enhancers and promoters, respectively.

A candidate distal enhancer (DE) bound by STAT5 and other TFs, including GR, NFIB and MED1, was identified 2.3 kb 5’ of the *Csn1s2b* TSS. STAT5 and NFIB binding coincided with their respective recognition motifs, suggesting their direct attachment to DNA. Unique to the *Csn1s2b* DE is the independent and physically separated binding of STAT5 and NFIB, which is not observed in the *Csn1s2a* locus (Fig. 1c-d), suggesting the possibility of distinct contributions in establishing a functional enhancer. Additional TF binding was detected within the first intron of the *Csn1s2b* gene (Fig. 1b). The presence of H3K4me1 marks in the DE regions affirms their enhancer status. TF binding at the intronic site was distinctly different from the upstream sites (Supplementary Fig. 1a). Both, TF binding and histone marks were strong during pregnancy and declined during lactation (Fig. 1b and Supplementary Fig. 1a), suggesting that this site might be necessary to activate the locus or suppressing gene expression during pregnancy.

Enhancers are occupied by Pol II which can result in the transcription of eRNAs^11^. However, their importance in enhancer-mediated gene activation remains elusive^12–14^. Since the *Csn1s2b* enhancer is occupied by Pol II (Fig. 1d) and activates gene expression more than 250-fold (Fig. 1a), we asked whether it is associated with eRNAs. We conducted total RNA-seq from mammary tissue and analyzed the presence of eRNAs in the *Csn1s2a* and *Csn1s2b* loci (Supplementary Fig. 2a-e). Despite the presence of Pol II, no eRNAs were detected at the *Csn1s2b* enhancer (Supplementary Fig. 2c). In contrast, eRNAs were detected at the *Csn1s2a* candidate enhancer and the *Csn1s2b* intronic candidate enhancer (Supplementary Fig. 2a and e). This suggests that eRNAs are not required for the activity of the *Csn1s2b* distal enhancer.

### Canonical TF motifs are not required for the establishment and function of the enhancer

Next, we conducted genetic experiments to determine the biological significance of the *Csn1s2b* DE. We addressed the potential function of the two canonical GAS motifs (TTCnnnGAA) recognized by STAT5 and the NFIB motif (TGGCA), all of which associate with the ChIP-seq peaks (Fig. 2a). A non-canonical GAS motif with a 4bp spacer (TTCnnnnGAA) was detected between the NFIB and GAS1 sites. We generated mice carrying individual or combinatorial deletions of the GAS and NFIB motifs (Fig. 2a). Deletion of the second STAT5 motif (ΔS2) had an insignificant impact on *Csn1s2b* expression (Fig. 2b), with STAT5 binding and H3K27ac reduced by ~40% (Fig. 2c-d and Supplementary Fig. 3a). This suggests a compensatory role of STAT5 site S1. The deletion of the NFIB motif did not affect *Csn1s2b* expression and NFIB binding was overtly unimpeded (Fig. 2b-c) suggesting that it functions indirectly through STAT5. Even the combined deletion of the NFIB and GAS2 sites (ΔN/S2) on *Csn1s2b* expression and TF binding was not significant (Fig. 2b-c), suggesting that one binding site within the complex enhancer might be sufficient for its function. The STAT5 and NFIB coverage in the *Csn12b* enhancer of mutants (ΔS2, ΔN and ΔN/S2) was confirmed by the raw read mapping (Supplementary Fig. 3b), demonstrating that STAT5 and NFIB still bind to the mutant enhancer. These results suggest that a single TF might be sufficient to anchor additional TFs and establish a complex enhancer structure.

**Fig. 2.**
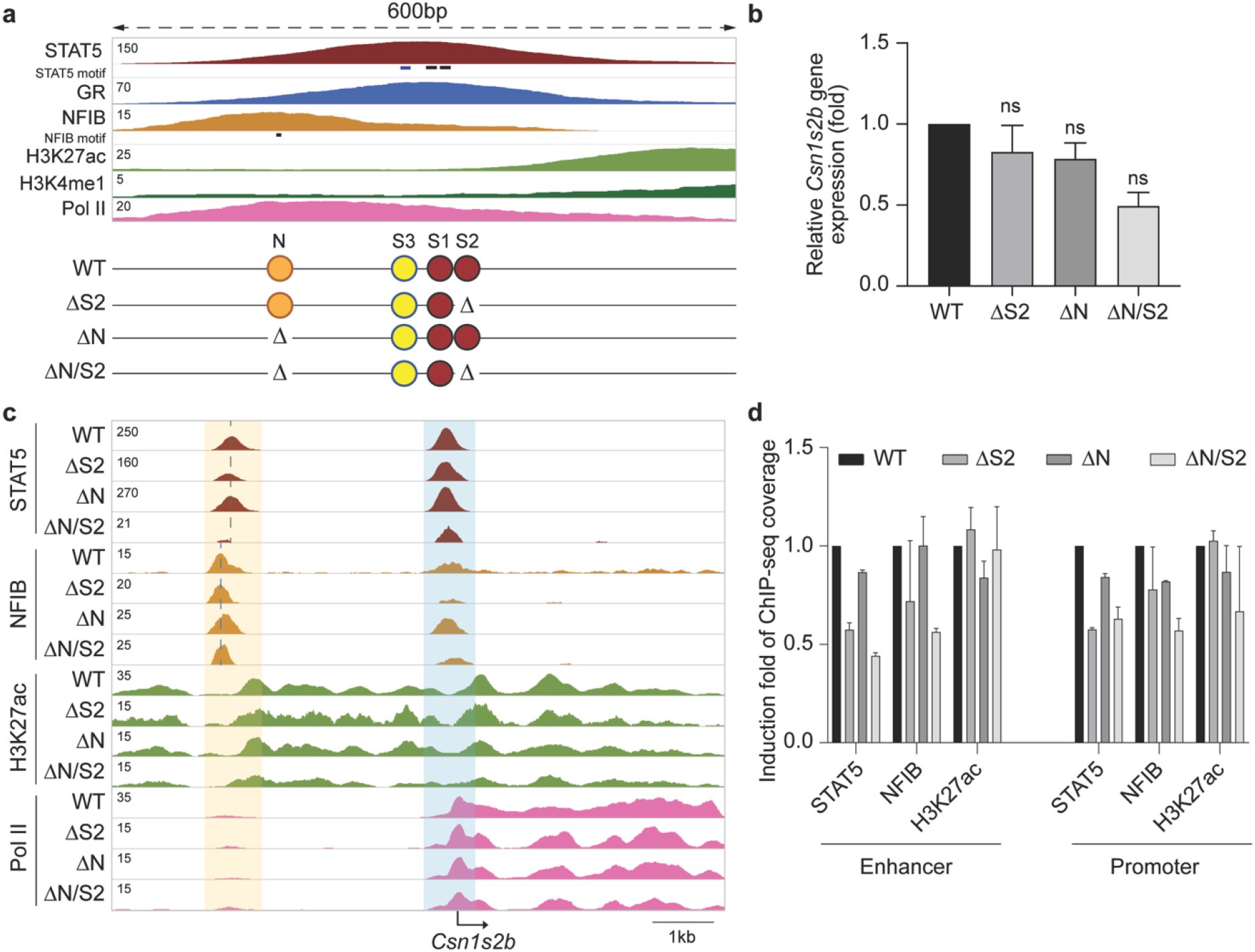
Limited function of STAT5 and NFIB sites in the *Csn1s2b* distal enhancer. **a** Genomic feature of the *Csn1s2b* distal enhancer and diagram of the deletions introduced in the mouse genome using CRISPR/Cas9 genome editing. TF binding sites were deleted individually (ΔS2 and ΔN) or in combination (ΔN/S2). While S1 and S2 display canonical GAS motifs (burgundy circles), S3 (yellow circle) has a non-canonical sequence. **b***Csn1s2b* mRNA levels in day 10 lactating (L10) mammary tissues from WT and mutant mice were measured by qRT–PCR and normalized to *Gapdh* levels. The *Cish* gene served as control. Results are shown as the means ± s.e.m. of independent biological replicates (WT and ΔN, *n* = 5; ΔS2 and ΔN/S2, *n* = 3). ANOVA was used to evaluate the statistical significance of differences between WT and mutant mice. ns, not significant. **c** The *Csn1s2b* locus was profiled using ChIP-seq data of WT and mutant tissue. **d** Reduction was calculated after variation between data set was normalized with *Cish* promoter coverage. Results are shown as the means ± s.e.m. of independent biological replicates (*n* = 2 to 4).

### A non-canonical GAS motif is critical for *Csn1s2b* gene expression during lactation

To address the possibility of additional TF binding sites, we dug deeper and analyzed the remaining sequences under the ChIP-seq peak, only to identify a non-canonical GAS motif with a 4 bp spacer (TTCnnnnGAA) (S3 in Fig. 3a). To explore the significance, we generated mice carrying S3 deletions in combination with NFIB and STAT5 sites that had little or no apparent biological activity by themselves (ΔN/S3, ΔN/S1/3 and ΔS2/3) (Fig. 3a). These deletions resulted in an 86% (ΔN/S3), 89% (ΔN/S1/3), and 96% (ΔS2/3) reduction of *Csn1s2b* mRNA (Fig. 3b), which was accompanied by a reduction of TF occupancy and H3K27ac marks (Fig. 3c-d and Supplementary Fig. 4). The combined absence of STAT5 sites S2 and S3 resulted not only in a complete absence of TF binding at the distal enhancer but also a sharp reduction at the promoter proximal site (Fig. 3c-d), in agreement with an almost complete abrogation of *Csn1s2b* expression. These results provide evidence that a non-canonical STAT5 binding motif is a key element in this enhancer and synergizes with a canonical site (S2). The integration of results from all the mutants strongly suggests that STAT5 preferentially binds at the non-canonical site S3.

**Fig. 3.**
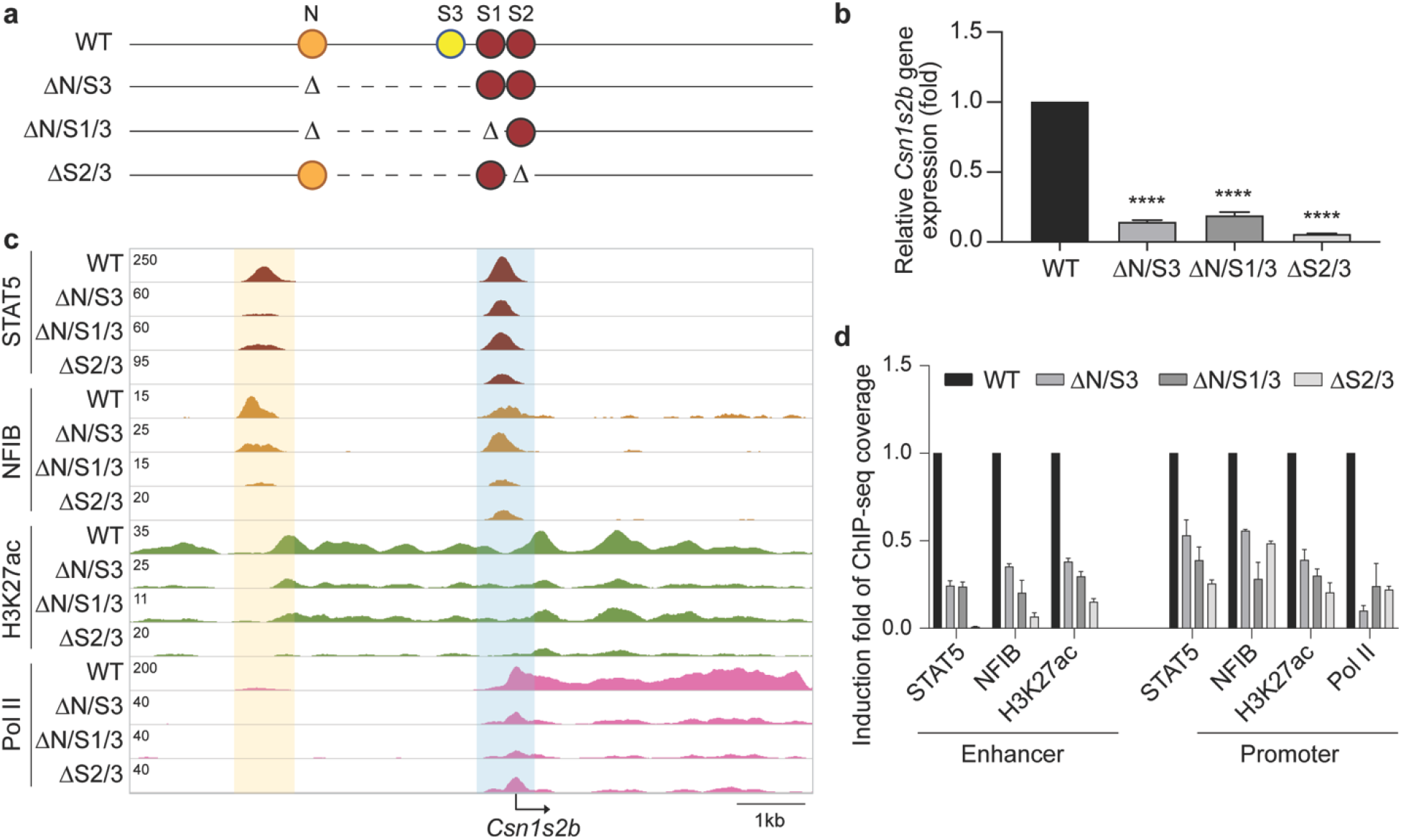
Requirement of a non-canonical STAT5 site in the *Csn1s2b* distal enhancer. **a** Diagram of the enhancer deletions introduced using CRISPR/Cas9 genome editing. The non-canonical STAT5 motif was deleted in combination with other enhancer motifs (ΔN/S3, ΔN/S1/3 and ΔS2/3). The canonical GAS motifs S1 and S2 are shown as burgundy circles, and the non-canonical GAS motif S3 is shown in yellow. **b***Csn1s2b* mRNA levels were measured by qRT-PCR in day 10 lactating (L10) mammary tissue of WT and mutant mice and normalized to *Gapdh* levels. The *Cish* gene was used as control. Results are shown as the means ± s.e.m. of independent biological replicates (WT, *n* = 5; ΔN/S3, ΔN/S1/3 and ΔS2/3, *n* = 4). ANOVA was used to evaluate the statistical significance of differences in WT and mutants. \*\*\*\**P* < 0.00001. **c** The consequences of enhancer deletions were confirmed by STAT5A, NFIB and H3K27ac ChIP-seq analysis in WT and mutant tissues at L10. The *Cish* locus served as control (Supplementary Fig. 3). **d** Reduction was calculated after variation between data set was normalized by *Cish* promoter coverage. Results are shown as the means ± s.e.m. of independent biological replicates (*n* = 2 to 4).

### Stage-restricted activity of a candidate *Csn1s2b* intronic enhancer

Our ChIP-seq data also revealed a putative intronic enhancer in the *Csn1s2b* gene body (Fig. 1b and Supplementary Fig. 1a). STAT5 binding to this site is present at p18 and L1 and disappears at L10 suggesting the possibility that this site is required to activate the *Csn1s2b* locus during pregnancy. To test this hypothesis, we deleted the GAS motif in the intronic enhancer (ΔIE) in the mouse genome (Fig. 4a). This resulted in a 50% reduction of *Csn1s2b* mRNA at L1 (Fig. 4b) suggesting that the intronic candidate enhancer might only modulate gene expression but is not essential. At L10 its impact was marginal suggesting the dominant role of the upstream enhancers.

**Fig. 4.**
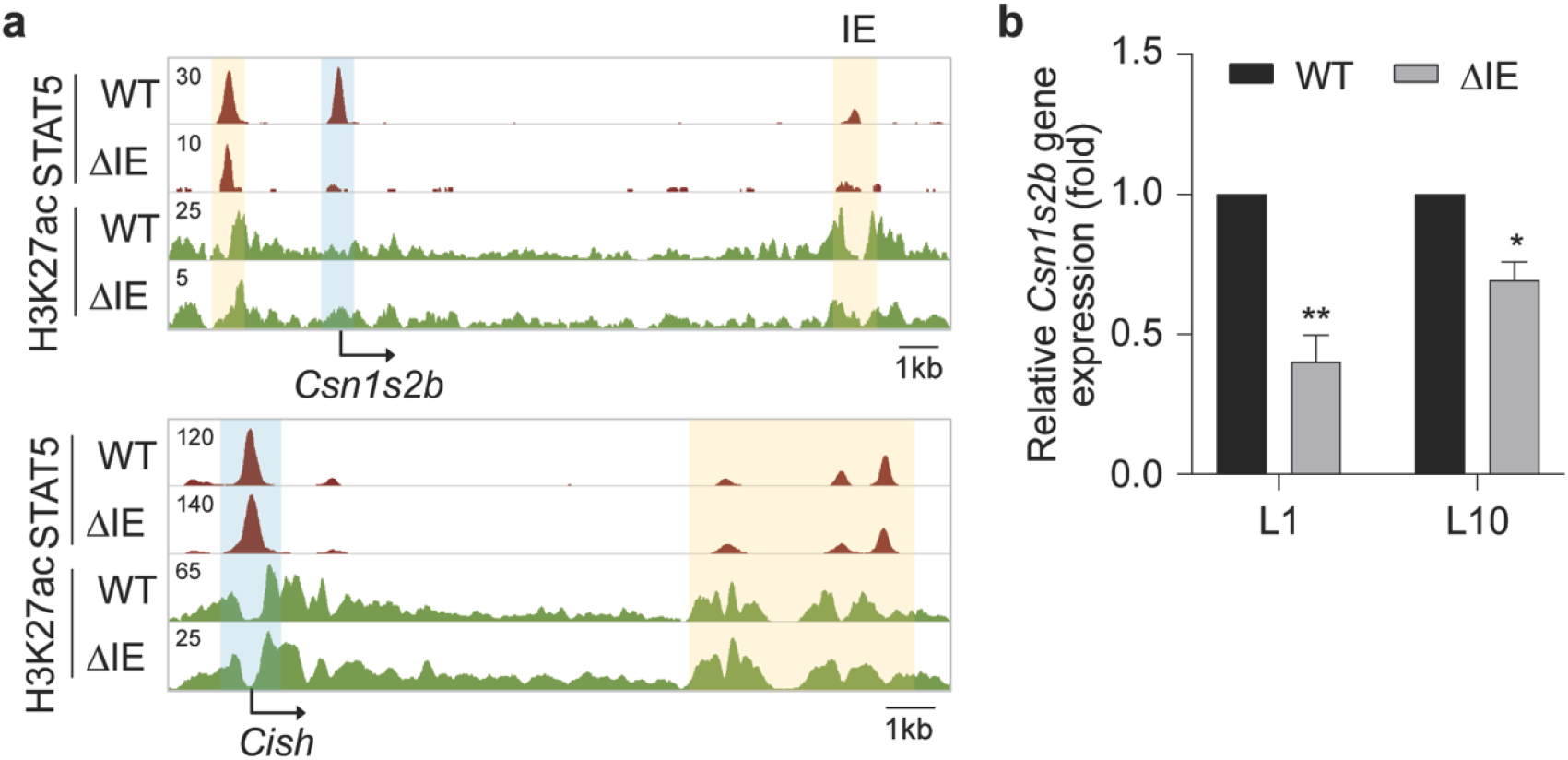
Limited contribution of the *Csn1s2b* intronic enhancer. **a** Genomic feature of the *Csn1s2b* locus in L1 mammary tissue of WT and ΔIE mice. The *Cish* locus served as a ChIP-seq control. IE, intronic enhancer. **b***Csn1s2b* mRNA levels were measured by qRT-PCR in L1 and L10 mammary tissue from WT and ΔIE mutant mice and normalized to *Gapdh* levels. The *Cish* gene was used as controls. Results are shown as the means ± s.e.m. of independent biological replicates (WT, *n* = 5; ΔIE, *n* = 4). *t*-test was used to evaluate the statistical significance of differences in WT and mutant. \**P* < 0.05, \*\**P* < 0.001.

## Discussion

Here we provide genetic clarity for the biological significance of building blocks within a complex enhancer responding to lactation hormones and driven by the collaboration of key lineage transcription factors. These analyses exposed the plasticity within enhancers and the hitherto underappreciated concept that TF binding to their DNA motif is not required for their function. Through this newfound knowledge, we gained major new insight into how a lineage-specific enhancer utilizes the NFIB and STAT5 transcription factors in activating its target gene during lactation.

In acquiring this knowledge, we learned that the *Csn1s2b* enhancer, which is preferentially active during lactation, is fundamentally different from other enhancers that activate genes in mammary tissue during pregnancy^15^. Although both enhancer classes use the cytokine-induced TF STAT5 as their core building block, their temporal recruitment is unique and the *Csn1s2b* enhancer is preferentially occupied during lactation. Mechanistically, we propose that this unique activation is the result of STAT5 binding to a non-canonical site. Our finding that NFIB, a critical co-activator for *Csn1s2b*^16^, does not depend on its DNA recognition motif adds further intrigue and provides evidence that the recruitment of multiple TFs can be facilitated through a single anchor, STAT5 in mammary enhancers.

Enhancers and super-enhancers can display additivity and synergy^17–20^, which is likely controlled by the co-binding of several TFs^15,19–21^. Deletion of individual TF sites within enhancers does not necessarily lead to its impairment^19,21^, suggesting additional complexity and compensation. While our previous studies have revealed that some mammary enhancers and super-enhancers feature a single TF binding site that anchors the key and non-redundant STAT5^15,22^, this work has revealed additional complexity within a uniquely strong enhancer. This complexity of TF binding might explain the stage-specificity and strength of this developmentally regulated enhancer. Most intriguingly, there was no absolute requirement for some of the individual binding motifs as plasticity permitted binding of individual TF in the absence of their motif through juxtaposed components.

The increasing use of a wide range of ChIP-seq and chromatin capture approaches suggest that the mammalian genome is riddled with candidate enhancers that control the spatio-temporal expression of lineage-specific genes^11,23–26^. However, the genuine biological significance of putative enhancers is far from clear and, as shown here, uncovering their function and complexity requires detailed genetic studies. Understanding the true biological significance of enhancers through experimentation is arduous and time consuming and a large number of candidate enhancers that had been predicted by genome-wide screens will fail functional tests. As future inquiries are conducted, the multiplicity of enhancer components activated by external inputs will continue to unfold.

## Methods

### Mice

All animals were housed and handled according to the Guide for the Care and Use of Laboratory Animals (8th edition) and all animal experiments were approved by the Animal Care and Use Committee (ACUC) of National Institute of Diabetes and Digestive and Kidney Diseases (NIDDK, MD) and performed under the NIDDK animal protocol K089-LGP-17. CRISPR/Cas9 targeted mice were generated using C57BL/6N mice (Charles River) by the transgenic core of the National Heart, Lung, and Blood Institute (NHLBI). Single-guide RNAs (sgRNA) were obtained from either OriGene (Rockville, MD) or Thermo Fisher Scientific (Supplementary Table 2). Target-specific sgRNAs and *in vitro* transcribed *Cas9* mRNA were co-microinjected into the cytoplasm of fertilized eggs for founder mouse production. The ΔN/S2 and ΔS2/3 mutant mouse was generated by injecting sgRNAs for NFIB site into zygotes collected from ΔS2 mutant mice. All mice were genotyped by PCR amplification and Sanger sequencing (Macrogen and Quintara Biosciences) with genomic DNA from mouse tails (Supplementary Table 3).

### Chromatin immunoprecipitation sequencing (ChIP-seq) and data analysis

Mammary tissues from specific stages during pregnancy and lactation were harvested, and stored at −80°C. The frozen-stored tissues were ground into powder in liquid nitrogen. Chromatin was fixed with formaldehyde (1% final concentration) for 15 min at room temperature, and then quenched with glycine (0.125 M final concentration). Samples were processed as previously described^22^. The following antibodies were used for ChIP-seq: STAT5A (Santa Cruz Biotechnology, sc-1081 and sc-271542), GR (Thermo Fisher Scientific, PA1-511A), NFIB (Sigma-Aldrich, HPA003956), MED1 (Bethyl Laboratory, A300-793A), H3K27ac (Abcam, ab4729), RNA polymerase II (Abcam, ab5408), H3K4me1 (Active Motif, 39297) and H3K4me3 (Millipore, 07-473). Libraries for next-generation sequencing were prepared and sequenced with a HiSeq 2500 or 3000 instrument (Illumina). Quality filtering and alignment of the raw reads was done using Trimmomatic^27^ (version 0.36) and Bowtie^28^ (version 1.1.2), with the parameter ‘-m 1’ to keep only uniquely mapped reads, using the reference genome mm10. Picard tools (Broad Institute. Picard, http://broadinstitute.github.io/picard/. 2016) was used to remove duplicates and subsequently, Homer^29^ (version 4.8.2) and deepTools^30^ (version 3.1.3) software was applied to generate bedGraph files, seperately. Integrative Genomics Viewer^31^ (version 2.3.81) was used for visualization. Coverage plots were generated using Homer^29^ software with the bedGraph from deepTools as input. R and the packages dplyr (https://CRAN.R-project.org/package=dplyr) and ggplot2^32^ were used for visualization. Each ChIP-seq experiment was conducted for two replicates. Sequence read numbers were calculated using Samtools^33^ software with sorted bam files. The correlation between the ChIP-seq replicates was computed using deepTools using Spearman correlation.

### RNA isolation and quantitative real-time PCR (qRT–PCR)

Total RNA was extracted from frozen mammary tissue of wild type and mutant mice using a homogenizer and the PureLink RNA Mini kit according to the manufacturer’s instructions (Thermo Fisher Scientific). Total RNA (1 μg) was reverse transcribed for 50 min at 50°C using 50 μM oligo dT and 2 μl of SuperScript III (Thermo Fisher Scientific) in a 20 μl reaction. Quantitative real-time PCR (qRT-PCR) was performed using TaqMan probes (*Csn1s2a*, Mm00839343_m1; *Csn1s2b*, Mm00839674_m1; mouse *Gapdh*, Mm99999915_g1, Thermo Fisher scientific) on the CFX384 Real-Time PCR Detection System (Bio-Rad) according to the manufacturer’s instructions. PCR conditions were 95°C for 30s, 95°C for 15s, and 60°C for 30s for 40 cycles. All reactions were done in triplicate and normalized to the housekeeping gene *Gapdh*. Relative differences in PCR results were calculated using the comparative cycle threshold (*C*_*T*_) method.

### Total RNA-seq analysis

Total RNA-seq reads were analyzed using Trimmomatic^27^ (version 0.36) to check read quality (with following parameters: LEADING: 3, TRAILING: 3, SLIDINGWINDOW: 4:20, MINLEN: 36). The alignment was performed in Bowtie aligner^28^ (version 1.1.2) using paired end mode.

### Statistical analyses

For comparison of samples, data were presented as standard deviation in each group and were evaluated with a *t-*test and 2-way ANOVA multiple comparisons using PRISM GraphPad. Statistical significance was obtained by comparing the measures from wild-type or control group, and each mutant group. A value of \**P* < 0.05, \*\**P* < 0.001, \*\*\**P* < 0.0001, \*\*\*\**P* < 0.00001 was considered statistically significant.

## Data availability

All data were obtained or uploaded to Gene Expression Omnibus (GEO). ChIP-seq data of wild-type tissue at L1 and L10 were obtained under GSE74826, GSE119657, GSE115370 and GSE127144. RNA-seq data for WT at p18, L1 and L10 were downloaded from GSE115370 and GSE127144. The ChIP-seq data for WT and mutant mice will be uploaded in GEO.

## Acknowledgements

We thank Ilhan Akan, Sijung Yun and Harold Smith from the NIDDK genomics core for NGS. This work utilized the computational resources of the NIH HPC Biowulf cluster (http://hpc.nih.gov).

This work was supported by the Intramural Research Programs (IRPs) of National Institute of Diabetes and Digestive and Kidney Diseases (NIDDK) and National Heart, Lung, and Blood Institute (NHLBI).

## Author contributions

H.K.L. and L.H. designed the study. C.L. generated mutant mice. H.K.L. established mutant mouse line and performed experiments, data analysis and computational analysis. H.K.L. and L.H. supervised the study and wrote the manuscript. All authors approved the final version.

## References

1. Ong, C.T. & Corces, V.G. Enhancer function: new insights into the regulation of tissue-specific gene expression. Nat Rev Genet 12, 283–93 (2011).

2. Andersson, R. & Sandelin, A. Determinants of enhancer and promoter activities of regulatory elements. Nat Rev Genet 21, 71–87 (2020).

3. Cui, Y. et al. Inactivation of Stat5 in mouse mammary epithelium during pregnancy reveals distinct functions in cell proliferation, survival, and differentiation. Mol Cell Biol 24, 8037–47 (2004).

4. Liu, X. et al. Stat5a is mandatory for adult mammary gland development and lactogenesis. Genes Dev 11, 179–86 (1997).

5. Shillingford, J.M. et al. Jak2 is an essential tyrosine kinase involved in pregnancy-mediated development of mammary secretory epithelium. Mol Endocrinol 16, 563–70 (2002).

6. Yamaji, D., Kang, K., Robinson, G.W. & Hennighausen, L. Sequential activation of genetic programs in mouse mammary epithelium during pregnancy depends on STAT5A/B concentration. Nucleic Acids Res 41, 1622–36 (2013).

7. Hennighausen, L.G. & Sippel, A.E. Characterization and cloning of the mRNAs specific for the lactating mouse mammary gland. Eur J Biochem 125, 131–41 (1982).

8. Lee, H.K., Willi, M., Shin, H.Y., Liu, C. & Hennighausen, L. Progressing super-enhancer landscape during mammary differentiation controls tissue-specific gene regulation. Nucleic Acids Res 46, 10796–10809 (2018).

9. Groenen, M.A., Dijkhof, R.J., Verstege, A.J. & van der Poel, J.J. The complete sequence of the gene encoding bovine alpha s2-casein. Gene 123, 187–93 (1993).

10. Rijnkels, M., Elnitski, L., Miller, W. & Rosen, J.M. Multispecies comparative analysis of a mammalian-specific genomic domain encoding secretory proteins. Genomics 82, 417–32 (2003).

11. Hirabayashi, S. et al. NET-CAGE characterizes the dynamics and topology of human transcribed cis-regulatory elements. Nat Genet 51, 1369–1379 (2019).

12. Catarino, R.R. & Stark, A. Assessing sufficiency and necessity of enhancer activities for gene expression and the mechanisms of transcription activation. Genes Dev 32, 202–223 (2018).

13. Lai, F., Gardini, A., Zhang, A. & Shiekhattar, R. Integrator mediates the biogenesis of enhancer RNAs. Nature 525, 399–403 (2015).

14. Willi, M. et al. Facultative CTCF sites moderate mammary super-enhancer activity and regulate juxtaposed gene in non-mammary cells. Nat Commun 8, 16069 (2017).

15. Shin, H.Y. et al. Hierarchy within the mammary STAT5-driven Wap super-enhancer. Nat Genet 48, 904–911 (2016).

16. Robinson, G.W. et al. Coregulation of genetic programs by the transcription factors NFIB and STAT5. Mol Endocrinol 28, 758–67 (2014).

17. Hay, D. et al. Genetic dissection of the alpha-globin super-enhancer in vivo. Nat Genet 48, 895–903 (2016).

18. Will, A.J. et al. Composition and dosage of a multipartite enhancer cluster control developmental expression of Ihh (Indian hedgehog). Nat Genet 49, 1539–1545 (2017).

19. Croft, B. et al. Human sex reversal is caused by duplication or deletion of core enhancers upstream of SOX9. Nat Commun 9, 5319 (2018).

20. Man, J.C.K. et al. An enhancer cluster controls gene activity and topology of the SCN5A-SCN10A locus in vivo. Nat Commun 10, 4943 (2019).

21. Okamura, E. et al. Esrrb function is required for proper primordial germ cell development in presomite stage mouse embryos. Dev Biol 455, 382–392 (2019).

22. Metser, G. et al. An autoregulatory enhancer controls mammary-specific STAT5 functions. Nucleic Acids Res 44, 1052–63 (2016).

23. Gaffney, D.J. Mapping and predicting gene-enhancer interactions. Nat Genet 51, 1662–1663 (2019).

24. Fulco, C.P. et al. Activity-by-contact model of enhancer-promoter regulation from thousands of CRISPR perturbations. Nat Genet 51, 1664–1669 (2019).

25. Chen, H. et al. Dynamic interplay between enhancer-promoter topology and gene activity. Nat Genet 50, 1296–1303 (2018).

26. Jung, I. et al. A compendium of promoter-centered long-range chromatin interactions in the human genome. Nat Genet 51, 1442–1449 (2019).

27. Bolger, A.M., Lohse, M. & Usadel, B. Trimmomatic: a flexible trimmer for Illumina sequence data. Bioinformatics 30, 2114–20 (2014).

28. Langmead, B., Trapnell, C., Pop, M. & Salzberg, S.L. Ultrafast and memory-efficient alignment of short DNA sequences to the human genome. Genome Biol 10, R25 (2009).

29. Heinz, S. et al. Simple combinations of lineage-determining transcription factors prime cis-regulatory elements required for macrophage and B cell identities. Mol Cell 38, 576–89 (2010).

30. Ramirez, F. et al. deepTools2: a next generation web server for deep-sequencing data analysis. Nucleic Acids Res 44, W160–5 (2016).

31. Thorvaldsdottir, H., Robinson, J.T. & Mesirov, J.P. Integrative Genomics Viewer (IGV): high-performance genomics data visualization and exploration. Brief Bioinform 14, 178–92 (2013).

32. Love, M.I., Huber, W. & Anders, S. Moderated estimation of fold change and dispersion for RNA-seq data with DESeq2. Genome Biol 15, 550 (2014).

33. Masella, A.P. et al. BAMQL: a query language for extracting reads from BAM files. BMC Bioinformatics 17, 305 (2016).

